# Histological evidences of diabetic kidney disease precede clinical diagnosis

**DOI:** 10.1101/493890

**Authors:** Giorgia Comai, Deborah Malvi, Andrea Angeletti, Francesco Vasuri, Sabrina Valente, Francesca Ambrosi, Irene Capelli, Matteo Ravaioli, Gianandrea Pasquinelli, Antonietta D’Errico, Alessia Fornoni, Gaetano La Manna

## Abstract

In the absence of a histhological diagnosis, persistent albuminuria is globally accepted as the main diagnostic criteria for diabetic kidney disease.

In the present retrospective study, we evaluated data from an Italian cohort of 42 diabetic deceased donors (mainly type 2 diabetes). Using kidney biopsies obtained at time of donation to evaluate single or double allocation based on Karpinski score, we determined the prevalence of histological lesions attributable to diabetes.

All 42 donors presented with normoalbuminuria and an estimated Glomerular Filtration Rate (CKD-EPI) > 60 ml/min/1.73m^2^. Of the 36 patients with available kidney biopsy, one was non interpretable and 32 had histopathological lesions consistent with diabetic kidney disease and encompassing all histological classes.

Thus, we found a relatively high proportion of histologically proven diabetic kidney disease that was clinically undiagnosed as none of the patient had significant albuminuria and eGFR <60 cc/min/1.73 m2. Our data support the need to implement routine kidney biopsies in normoalbuminuric diabetic subjects in early stages of Chronic Kidney Disease. As others have shown that histological lesions associated with diabetic kidney disease predict disease progression, such strategy may help to risk stratify patients and may guide therapeutic decisions at early disease stages.

## Introduction

Chronic kidney disease (CKD) is a worldwide public health problem, and represents one of the principal causes of cardiovascular mortality around the world (1). Diabetic kidney disease (DKD), which develops in approximately 10% to 30% of patients with diabetes mellitus (2), is one of the most important long-term complications of diabetes and it is among the leading causes of end stage kidney disease (ESKD) in developed countries; the overlap of diabetes mellitus and renal injury exponentially amplifies the death risk and cardiovascular events (3). DKD is characterized by the development of albuminuria often followed over a long period of time (10-20 years) by a decline in glomerular filtration rate (GFR) towards ESRD (4–6). Peculiar histological features that typically characterize DKD are glomerular basement membrane thickening, nodular or diffuse mesangial sclerosis, arteriolar hyalinosis, micro aneurysms and exudative lesions (7). However, increasing evidence indicates that many patients with diabetes erroneously labelled as having progressive forms of DKD are rather developing non-diabetic renal diseases (NDRD) or ‘mixed’ conditions where typical features of DKD overlap with other pathological diagnosis (7–11). Therefore, while clinically indicated kidney biopsies in DKD are likely to reveal NDRD, routine kidney biopsies are likely to provide a better understanding of the natural history of DKD. This was clearly demonstrated by studies that implemented research kidney biopsies at early disease stage. A unique feature of DKD, thickening of the glomerular basement membrane predict DKD progression in a noromoalbuminuric cohort of patients with type 1 diabetes (T1D) (12), while a reduction in podocytes number is observed at later stages (13). In type 2 diabetes (T2D), several histological lesions including podocytes number precede albuminuria (14–16). Furthermore, lesions in T2D are rather heterogeneous, and only typical diabetic nephropathy features were described to contribute to loss of GFR (17). The correct classification of such patients would be crucial to predict the natural course and early treat the disease, thus allowing the appropriate therapeutic treatment in a timely manner. The classic paradigm, representing the natural history of DKD would expect an early phase of microalbuminuria (albumin excretion of more than 30 mg/g of creatinine in 2 out of 3 random urine samples collected in within a 6 months period), which precedes overt nephropathy, characterized by macroalbuminuria and a progressive decline of GFR (18). Despite such a large body of evidence of how histological predictor can predict DKD progression, current guidelines for screening for DKD are solely based on the detection of persistent albuminuria in two out of three morning urine collections over a six months period of time (19). This is a major limitation, as many diabetic patients who reach a persistent GFR<60 cc/min remain normoalbuminuric (20–24). Therefore the definition of DKD that recognizes the clinical appearance of albuminuria as specific of microangiopathic damage may not be so exhaustive (8, 25).

With this retrospective study we collected and analyzed data from a cohort of patients with positive history for diabetes, enrolled as candidate cadaveric donors for kidney transplant. The aim of the present study was to investigate the prevalence and classification of histopathological lesions of DKD in diabetic patients with clinically silent nephropathy.

## Methods

We performed a retrospective inclusion of diabetic patients considered suitable as deceased donors for kidney transplant, by 2004 to 2015 at S. Orsola Hospital-University of Bologna, Italy. At time of donation, kidneys from diabetic patients were biopsied to evaluate single or double allocation based on Karpinski score (26), in accordance to Italian Clinical Guidelines for Organ Transplantation from Deceased Donors published by Italian National Transplant Centre (CNT)-Italian National Institute of Health (ISS) (http://www.trapianti.salute.gov.it). We considered both T1D and T2D in candidate donors who were older than 18 years and with available kidney biopsy at time of death.

### Clinical data

Clinical informations were obtained through the medical records available at death of patients by Intensive Care Unites where patients were hospitalized and provided to the Regional Transplant Center of Emilia Romagna in Bologna once becoming eligible as donors. Due to the retrospective nature of this study, registration or approval by the ethics committee was waived. The study was carried out in conformity with the Declaration of Helsinki. The following laboratory and clinical measurements were included: serum creatinine at time of hospitalization, eGFR (calculated using the CKD-EPI formula), albuminuria (> 30 mg/dl) determined by spot urine collections, serum hemoglobin and serum cholesterol. Data that reflected a stable representation of the serum and urine levels were included. We also obtained data regarding comorbidities, medication history, hypertension, smoking, and complications secondary to diabetes. Causes of death were obtained from the hospitalization records.

### Histopathological analysis

Renal tissue was fixed in Serra solution and embedded in paraffin. Slices were cut at 3- μm thickness and stained with haematoxylin and eosin, periodic acid–Schiff (PAS) and Masson’s trichrome stain. Renal tissue specimens were scored by 2 pathologists blinded each other and to the patients’ clinical data. Glomerular lesions, interstitial lesions, and vascular lesions were scored in accordance with the established histopathologic classification for DKD. Diabetes score were assessed according to Tervaert *et al*. (10, 27): patients without histologic lesions, and with no thickening of the glomerular basal membrane at Transmission Electron Microscopy (TEM), were designated as class 0 DKD; patients without histologic lesions, but with thickening of the glomerular basal membrane (>430 nm in males and >395 nm in females) at TEM, were designated as class I DKD; patients showing mild mesangial expansion were designated as class IIa DKD; patients showing moderate/severe mesangial expansion were designated as class IIb DKD; patients with patent diabetic glomerular nodules and <50% globally sclerotic glomeruli were designated as class III DKD; patients with ≥50% globally sclerotic glomeruli were designated as class IV DKD. The four histological parameters of the Karpiski score (26) were separately assessed as well: glomerulosclerosis score, tubular atrophy score, interstitial fibrosis and vascular damage (**Supplemental Data**). Finally, further histopathological variables were evaluated for each kidney as reported in detailed in **Supplemental Data**.

### Transmission Electron Microscopy

In the present study, TEM was applied in order to measure the thickness of the glomerular basal membrane and distinguish class 0 from class I DKD. Small specimens of renal tissue were retrieved from paraffin blocks, de-paraffined in xylene, rehydrated in ethanol (100%, 95% and 70%) and washed in 0.15 M sodium cacodylate buffer. After post-fixation in 1% osmium tetroxide, the samples were with increasing concentration of ethanol (from 70 % to 100%), embedded in Araldite resin and cut with the ultramicrotome. Ultrathin sections were stained with uranyl acetate and lead citrate before the examination with Philips CM10 (FEI Company, Milan, Italy) Transmission Electron Microscope equipped with a Gatan camera. For each sample, five digital images were randomly acquired using the FEI proprietary software Olympus SIS Megaview SSD digital camera. The thickness of the Glomerular Basement Membrane (GBM) was measured in twelve different positions at 13500 of magnification. Class I DKD was defined as the presence of basal membranes with average thickness >430 nm in male patients and >395 nm in female patients (28).

### Statistical analysis

Differences with a P value less than 0.05 were considered statistically significant. The histopathological data were analyzed using the chi-square test.

## Results

A total of 42 cadaveric kidney donors were selected based on established diagnosis of type 1 or type 2 diabetes prior to expiration. Of those, 7 were excluded for lack of available kidney biopsy or poor quality of the kidney biopsy. Therefore 35 diabetic subjects were included in the analysis. The characteristics of the 35 patients are summarized in Table 1 Individual data are also provided in **S1 Table**. The cohort was characterized by 1 patients with T1D and 34 patients with T2D. The mean age was 69.7 years and 51.4% of subjects were female.

### Clinical data

4 of 35 patients were treated with insulin, 14 patients received oral anti-diabetic agents while 20% (7 patients) achieved glycaemic control through lifestyle changes, mostly diet. Multiple dipstick testing for albuminuria available prior to expiration excluded the presence of albuminuria in all patients. Ultrasonography examination reported normal kidney size (11.63 cm ± 0.15). The medical records revealed that 24 patients (68.6%) had a diagnosis of primary hypertension, and 10 of them had received therapy with an angiotensin-converting enzyme inhibitor or an angiotensin-receptor blocker. Non of the patient had diabetic retinopathy (Table 1). Hemorrhagic stroke was the most prevalent cause of death, followed by ischemic stroke and cranial trauma (**Supplemental Table**).

**Table1.** Baseline characteristics of the patients

| Baseline characteristics | Mean ± SEM (%) |
| --- | --- |
| Sex (M/F) | 17/18 |
| Age (yrs) | $69.7 \pm 1.3$ |
| T1D/T2D | 1/34 |
| Proteinuria (mg/dl) | $11.02 \pm 3.2$ |
| Creatinine (mg/dl) | $0.98 \pm 0.07$ |
| CKD Class | Class I: 7 (20)<br>Class II: 28 (80) |
| Retinopathy | 0 |
| Hypertension | 24 (68.6) |
| Dislipidemia | 21 (60) |
| Renal Longitudinal Size (cm) | $11.63 \pm 0.15$ |
| Diabetic Therapy | Insulin: 4 (11.4)<br>OAD: 14 (40)<br>LC: 6 (17.1)<br>N/A: 11 (31.4) |
| Anti-Hypertensive Therapy | 18 (51.4) |
| RAASi | 8 (22.8) |

eGFR, estimated glomerular filtration rate; LC, lifestyle changes; N/A, not available; OAD, oral anti-diabetic; RAASi, renin angiotensin aldosterone system inhibitor; SEM, standard error mean; T1D, type 1 diabetes; T2D, type 2 diabetes.; yrs, years.

### Histological lesions

Histopathological characteristics are summarized in Table 2. Histopathological classification revealed that 6 patients had mild light microscopic changes: in these 6 cases, TEM was applied in order to distinguish class 0 from class I diabetic nephropathy(10). The mean glomerular basal membrane (GBM) thickness obtained was 260.566 ± 122.276 nm. Finally in 3 cases class 0 DKD was assigned, due to an average GBM thickness under the cut-off (see Methods section), while 3 cases were class I DKD. 22 patients were class IIa DKD, 3 patients class IIb DKD and 4 patients class III DKD (Figure 1); no class IV were observed in our series. Myointimal hyperplasia was observed in 45.7% patients, multifocal arteriolar hyalinosis in 60% of cases, Interstitial fibrosis and tubular atrophy (IFTA) in 62.8% of cases (**S1 Table**). NDRD lesions compatible with primary glomerulonephropathies were not reported.

**Table2.**
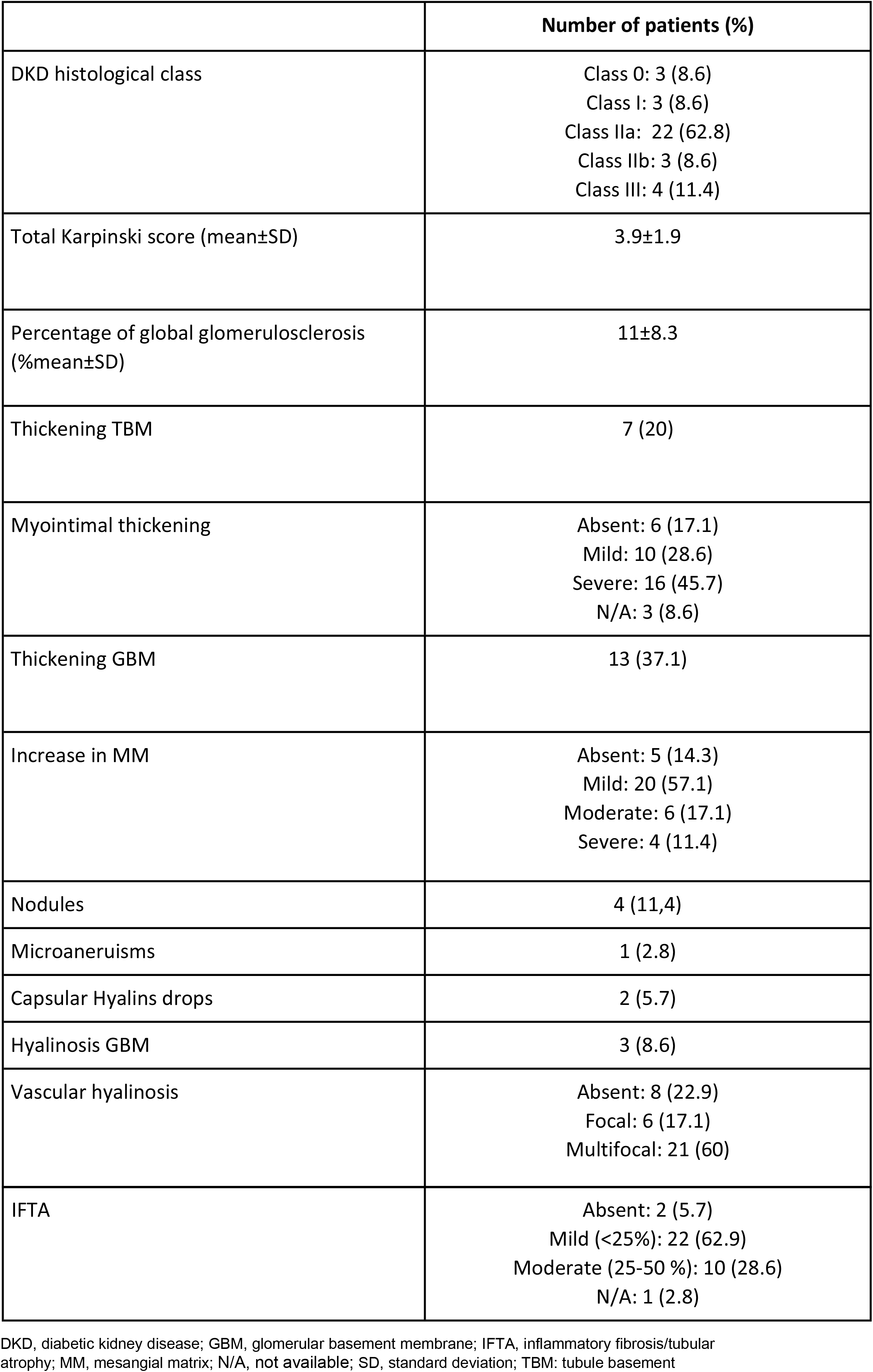
Histological findings

**Figure 1.**
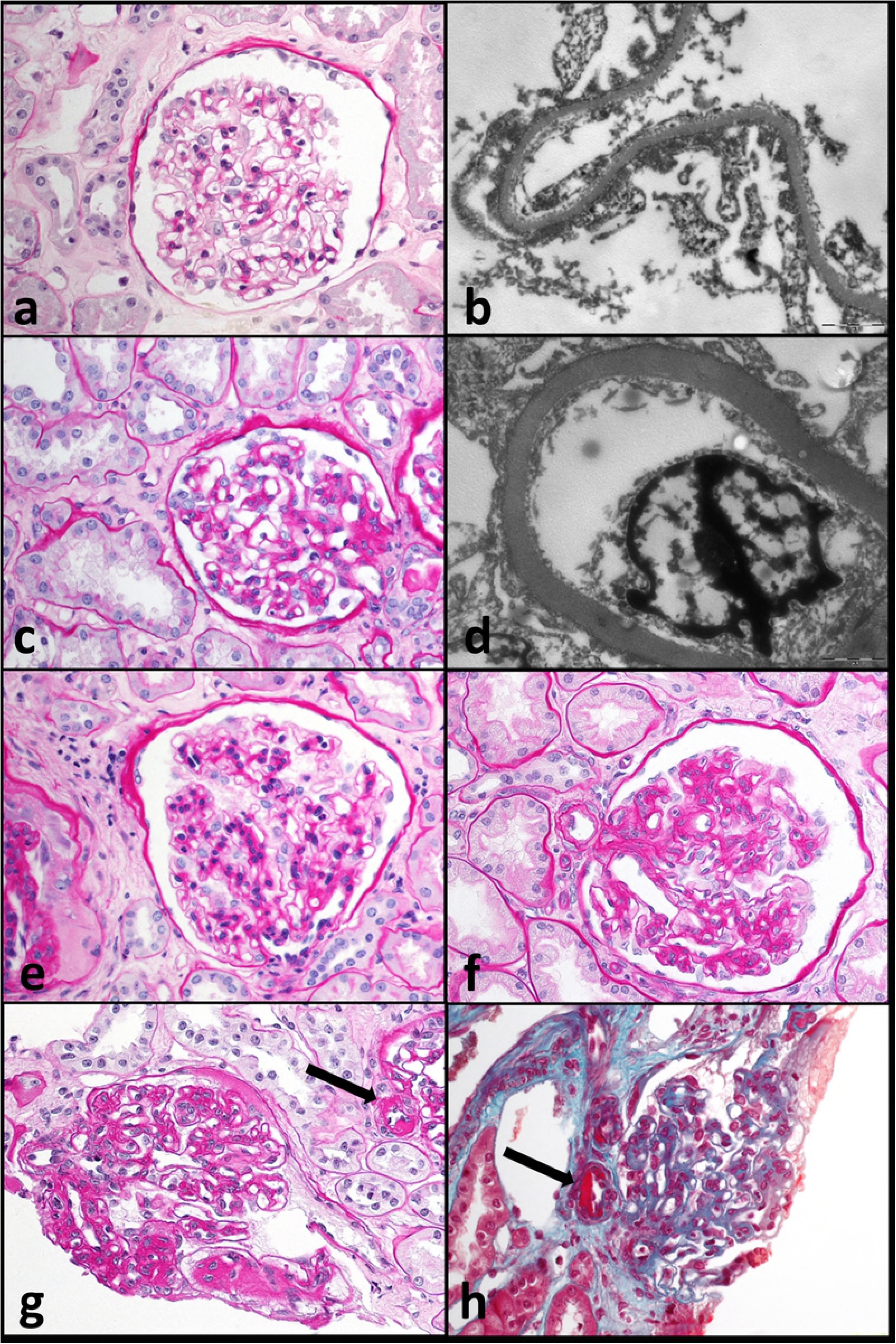
Examples of diabetic kidney disease classes at histology. Class 0 is characterized by normal-appearance glomeruli (a), without thickening of the glomerular basement membrane at transmission electron microscopy (b). Class I is characterized by a variable thickening of the glomerular basement membrane (c), requiring confirmation at transmission electron microscopy (d). Class II is characterized by mild (class IIa, square e) or moderate/severe (class IIb, square f) expansion of the glomerular mesangium. Class III is characterized by the presence of severe mesangial expansion, increased mesangial cellularity and Kimmelstiel-Wilson nodules (g); note the arteriolar hyalinosis (arrows). PAS (a,c,e,f,g) and Trichrome (h) stains; magnification 40x. Magnification 13500x (b,d).

## Discussion

Premised that absence of clinical signs or laboratory findings for DKD have represented a primary step to obtain eligibility as donors for kidney transplant, in our diabetic cohort of 35 suitable candidates as cadaveric donors for kidney transplant, histopathological changes compatible with DKD were present in 32 subjects. Differently to what the classic paradigm of DKD proposes (18), but consistent with prior natural history studies with research kidney biopsies in normoalbuminuric diabetic patients, our findings suggest that renal lesions consistent with DKD often develop before clinical diagnosis. Moreover, our data indicate that underdiagnosed DKD encompasses all DKD histological classes, including class III typically characterized by the presence of Kimmelstiel-Wilson nodules. The classic paradigm of DKD involves progressive steps from glomerular hyperfiltration, to microalbuminuria, to proteinuria and to lower GFR (18, 29). However in the last decade, this concept has been progressively changed with studies reporting a more heterogeneous presentation of DKD (9, 11), as example T2D patients with reduced eGFR presenting with no albuminuria (22, 30–32). In the DEMAND (Developing Education on Microalbuminuria for Awareness of Renal and Cardiovascular Risk in Diabetes) study, a cross-sectional study involving 11,573 patients with type 2 diabetes, 6072 individuals presented with normoalbuminuria and among them 17% (1,044 patients) had eGFR<60 ml/min per 1.73 m^2^ (32). Previously, Kramer *at al*. (30), in a cross-sectional study, reported this number being as high as 33%. These data are consistent with a recent report by Afkarian *et al*. (33), describing how prevalence of CKD (eGFR<60 ml/min per 1.73 m^2^), in patients with T2D, increased from 9.2% in 1988-1994 to 14.1% in 2009-2014, despite albuminuria decreased from 21% to 16% considering the same periods. However, a limited number of studies have previously described the presence of histologically proven DKD in absence of albuminuria and CKD. While most of these studies were done on prospective cohort of patients with research kidney biopsies, limited studies have validated these findings utilizing organs from cadaveric donors. Among them, in an autopsy study on 168 subjects with diabetes, Klessens *et al*. (34) have recently described 106 subjects with histopathologic changes compatible with DKD and 20 of the 106 did not present with DKD-associated proteinuria within their lifetime. Differently from our findings, half of patients considered by Klessens and colleagues were in CKD stage 3 or 4 (eGFR<60 ml/min per 1.73 m^2^) (34).

The presence of histologically proven DKD in our cohort is not associated with significant albuminuria. We speculate that the capability of healthy tubular epithelium to reabsorb proteins from the glomerular filtrate may hide glomerular protein leakage (35, 36). Only after interstitial damage occurred, the resulting loss of reabsorption capacity causes the albuminuria, providing a reasonable explanation why glomerular damage consistent with DKD can occur before the onset of albuminuria. Moreover, in some cases renin angiotensin aldosterone inhibitors administration may have delayed the onset of proteinuria (37).

Zhang *et al*. (38), in a cohort of 298 T2D patients, investigated clinical predictive characteristics of subclinical DKD: increased kidney size, abnormal levels of tubular injury markers, high blood pressure, and abnormal circadian rhythms were positive correlated with further development of DKD. Conversely, in our retrospective study, kidney size (corrected for Body Mass Index) and high blood pressure were not associated with histological severity of DKD.

Histologically, we assessed that the more represented DKD lesion was the class IIa, that means mild mesangial expansion. It must be kept in mind that our population is represented by marginal donors, with a mean age of 70 years. This justifies the nondiabetic alterations present, such as the mean total Karpinsky’s score (3.9) and the presence of IFTA in nearly 90%. As far as the vascular changes are concerned, a severe myointimal thickening was observed in 44.4% patients and a multifocal arteriolar hyalinosis in 58.3% of cases. These findings are not in contrast with previous data estimating around 80% the prevalence of not diabetic renal disease detected in renal biopsies of diabetic patients (39–41). However, previous studies may suffer of a selection bias regarding the reasons for performing renal biopsy (34): in our study, due to the different technique in providing renal biopsy, we had the huge benefit to study at least 30 glomeruli for most patient, which are certainly much more than glomeruli usually available.

Considering the amount of renal tissue, we are certain about the presence or absence of DKD, and, moreover, we are largely confident of its distribution over the DKD classes. Typical nodular lesions, were focally concentrated in specific areas of tissue. Thus, it is plausible that the not uniformed distribution may be partly responsible for the underpresentation of DKD in previously renal biopsy studies.

The early detection of histological DKD lesions in our population are consistent with reports by Fioretto *et al*. (42) and Caramori *et al*. (12), who elegantly evaluated the histological characteristics of DKD in patients with T1D, reporting how the width of the glomerular basement membrane predicts development of proteinuria and/or ESKD earlier than microalbuminuria, which is still largely accepted as the first clinical sign of DKD. Similarly to Klessens and colleagues (34), in our population the percentage of glomeruli with nodular mesangial sclerosis was very low (average value of 11 % in kidneys of patients in DKD class III) (**Supplemental Table**). Therefore, these data suggest that DKD histological lesions develop focally, in different moments, and that proteinuria may not develop until a certain percentage of glomeruli are not damaged. As possible mechanism, Said and Nasr (43) have suggested that in early DKD phases the tubules are not damaged and still able to reabsorb the leaked albumin from the injured glomeruli, contributing to the absence of albuminuria.

Currently, no standardized criteria for kidney biopsy in diabetic patients are reported and the decision to perform it is responsible of the physician alone. Until now, proteinuria with no proven diabetic retinopathy is the major recognized criteria for kidney biopsy (7, 44), in the present study we have shown as the histological classes of DKD are not directly associated with proteinuria, proving the kidney biopsy as the gold standard for early diagnosis of DKD.

Despite the advantages previously discussed, some limitations request a discussion. Proteinuria was detected by dipstick, therefore possible microalbuminuria was not detected. Moreover, the cohort may suffer from selection bias: population was totally composed of white individuals, therefore, the extension of these results to other populations is forbidden. Nodular lesions, characterizing for class III DKD may also occur in patients with hypertension and a history of smoking but without diabetes (especially in elderly subjects) (45). However, It seems unlikely that our results were strongly influenced by this entity, because patients with nodules had not an elderly median age (68 years old) and only 2 of 4 patients were in pharmacological treatment for hypertension.

In conclusion, our results show as advanced DKD histological lesions are present also with no significative proteinuria and in early stages of CKD, such as Class I and II. It is reasonable that when increase of albuminuria occurs, kidneys of diabetic patients may already manifest by minor to severe glomerular injuries. TEM results essential in the early stages of renal involvement in diabetic patient. Early identification of this underdiagnosed group of DKD is crucial to promptly provide a specific therapeutic regimen able to slow disease progression (3, 46, 47); the coming of new antidiabetic drugs enforces the needing (48). We acknowledge that a standardized kidney biopsy for all diabetic patients is logistically demanding, however, in consideration of our findings, the challenge may be performing kidney biopsy in diabetic subject at least with CKD Class II. Further studies should be designed to better investigate the development of diagnostic algorithms to accurately detect currently underdiagnosed patients.

## Acknowledgments

AF is supported by the NIH grants R01DK117599, R01DK104753, R01CA227493, U54DK083912, UM1DK100846, U01DK116101 and UL1TR000460 (Miami Clinical Translational Science Institute).

## Supporting information

S1 Table.

